# Comparison between Q_ST_ and Φ_ST_ indices in an endangered *Boswellia serrata* Roxb: Implications for conservation

**DOI:** 10.1101/442723

**Authors:** Shashank Mahesh, Pramod Kumar, Vivek Vaishnav, Naseer Mohammad, Shamim Akhtar Ansari

**Affiliations:** Tropical Forest Research Institute, PO-R.F.R.C., Mandla Road, Jabalpur-482 021, M.P., India; CSIR-National Botanical Research Institute, Lucknow-226001, Uttar Pradesh, India; Institute of Forest Productivity, Lalgutuwa, NH 23, Ranchi-835303, Jharkhand, India

**Keywords:** Location wise neutral marker trait index (Φ_ST_ (L)), Location wise quantitative trait index (Q_ST_ (L)), Location wise neutral marker trait index (Φ_ST_ (L)), Measures of population differentiation, Metapopulation wise quantitative trait index (Q_ST_ (P))

## Abstract

*Boswellia serrata*, an economically important indigenous tree of dry deciduous forests, provides oleoresin gum of pharmaceutical significance and excellent pulp for paper industries, but faces threat to extinction due to poor natural regeneration and commercial exploitation. 240 individuals of the species representing 12 locations of its natural distribution in central India were investigated to compare the genetic differentiation indices, Q_ST_ for GBH and wood fiber length and ϕ_ST_ for neutral (RAPD+ISSR) markers. The comparison for paired locations was more informative than for metapopulation. The most paired locations were either under the stabilizing selection (Q_ST_ (L) < Φ_ST_ (L)) or in the genetic drift (Q_ST_(L) = Φ_ST_ (L)) whereas a relatively small number of paired locations was under the divergent selection (Q_S T_(L) > Φ_ST_ (L)). The comparison for the metapopulation generating only a single trend of Q_ST_ (P) > Φ_ST_ (P) is, therefore, misleading. For conservation, the genetically deficit locations (Q_ST_ (L) < Φ_ST_ (L) and Q_ST_ (L) = Φ_ST_ (L)) of *B. serrata* warrant for reinforcement of their genetic diversity by introduction of genotypes from other genetically divergent locations (Q_ST_ (L) > Φ_ST_ (L)), which would check the fragmentation and genetic drift, resulting in reproductive vigour, natural regeneration and reverse the endangered status of the species.

## Introduction

A species sustains itself through diverse populations in space and time. These populations owe their origin from a genetically homogeneous stock in the antiquity that has assiduously experienced differentiation and selection through intrinsic genetic changes such as recombination, mutation, migration, etc. and adaptability to a specific set of climatic conditions, respectively. This implies that a locus experiences either selection or homeostasis, leading to its adaptability or neutrality, respectively. The loci governing quantitative traits are often subjected to selection pressure. The other intermittent loci do not experience selection pressure and remain neutral. The deviation of quantitative trait loci of under selection from neutral marker loci brings out the dynamics and adaptability at population or species levels. To judge it, two indices dealing with neutral trait loci and quantitative trait loci under selection have been devised as F_ST_ (Wright 1951) and Q_ST_ (Spitze 1993), respectively. A comparison between both indices provides three scenarios, *viz.*, Q_ST_ = F_ST_, Q_ST_ > F_ST_ and Q_ST_ < F_ST_. According to Merila and Crnokrak (2001), Q_ST_ = F_ST_ denotes degree of differentiation in quantitative traits due to genetic drift alone, Q_ST_ > F_ST_ means directional selection favouring dissimilar genotypes in different populations and Q_ST_ < F_ST_ exhibits retention of same genotypes across all populations. However, F_ST_ estimation tends to depreciate with the rate of mutation exceeding the migration rate within a population that may lead to upwardly biased comparison between both indices and erroneous inferences thereof (Edelaar et al. 2011). The difficulty may be surmounted by comparing Q_ST_ with Φ_ST_ (an F_ST_ analogue) of AMOVA (Excoffier and Lischer 2010). The Φ_ST_ is derived from genetic distance based allelic variations and makes neutral divergence independent of mutation rate (Edelaar et al. 2011). Further, the co-dominant SSR markers are prone to mutation rate as high as 10^-2^ (Ellegren 2004) and are not much suitable for estimation of F_ST_ for such comparison (Edelaar et al. 2011). In natural populations, the phenotypic variations are inseparable from the genetic and environmental contributions due to employment of purely phenotypic data of wild individuals for Q_ST_ computation. However, this constraint does not influence the outcome of the comparison between both indices (Leinonen et al. 2008).

*Boswellia serrata* Roxb. (family: Burseraceae), salai an endangered plant species is endemic to India and naturally distributed to central Indian states *viz.* Chhattisgarh, Madhya Pradesh, Parts of Rajasthan adjoining to Madhya Pradesh, Maharashtra and Orissa. The species thrives in very dry teak forests or in dry mixed deciduous forests in association with *Sterculia urens, Holarrhena pubescens* and *Anogeissus latifoila.* It is also an excellent source of pulp for production of very high quality paper. Rosin and turpentine from its oleoresin gum is used for manufacture of soap, paints, varnishes, printing ink, calico prints, etc (http://www.worldagroforestry.org). The gum fractions obtained from *B. serrata* have been found to be pharmaceutically very important as anti-oxidant (Assimopoulou et al. 2005), anti-inflammatory (Gayathari et al. 2007) and vaccine adjuvant (Khajuria et al. 2007).

However, *B. serrata* has been over exploited and exhibits scarce regeneration due to very poor seed set and germination arising from post-fertilization abortiveness of seeds, leading to fruits without seeds (Luna 1996; Sunnichan 2005). It is also classified as difficult to root species (Singh et al. 1992), but moderately responds to air layering (Singh et al. 2004). As a result, *B. serrata* has been declared as an endangered species with dwindling natural populations (Sharma 1983; Purohit et al. 1995; Ghorpade et al. 2010). An endangered species exhibits declining number and size of populations with impaired natural regeneration and inverted pyramid shaped age structure, i.e. post-reproductive individuals exceeding to pre-reproductive individuals (Jin 1997, 1998; Jin and Ke 2002). *B. serrata* shares several of these characteristics as it is represented by a few sporadic small populations restricted only in Madhya Pradesh and adjoining states.

Considering above in view, we conducted experiment to investigate the types of selection pressure and/or genetic drift enduring on endangered *B. serrata* metapopulation by comparing the quantitative genetic divergence (*Q_ST_*) of wood fiber length and GBH with neutral genetic divergence (*Φ_ST_*) of ISSR and RAPD markers.

## Materials and methods

### Plant sample

The naturally distributed metapopulation of the species was surveyed in central Indian state, Madhya Pradesh. Twelve locations representing eight different agroclimatic zones (Table 1 and Fig. 1) were sampled for wood radial cores and leaves. The samples were collected from 20 trees (separated by a distance of ≈ 100m apart from one and the other) of each location. Wood radial cores were extracted, and girth (GBH) was measured, at the breast height (1.34 m).

**Table 1.** Geo-climatic description of 12 collected locations of *Boswellia serrata* Roxb.

| Location | Code | GPS characteristics |  |  | Range |  |
| --- | --- | --- | --- | --- | --- | --- |
|  |  | Latitude (<sup>0</sup>N) | Longitude (<sup>0</sup>E) | Altitude (m) | Temperature (<sup>0</sup>C) | Rainfall (mm) |
| Balaghat | BG | 22.13 | 80.11 | 361.95 | 26-28 | 1500-1700 |
| Chhatarpur | CP | 24.65 | 79.79 | 284.55 | 36-38 | 1000-1100 |
| Chhindwara | CW | 22.42 | 78.63 | 635.35 | 26-28 | 1300-1400 |
| Damoh | DM | 23.30 | 79.25 | 454.30 | 28-30 | 1200-1300 |
| Gwalior | GW | 26.07 | 77.99 | 356.10 | 36-38 | 1000-1100 |
| Jabalpur | JB | 23.18 | 80.03 | 468.15 | 36-38 | 1400-1500 |
| Khandwa | KN | 21.48 | 76.29 | 468.70 | 26-28 | 1200-1300 |
| Mandla | MN | 22.90 | 80.20 | 555.40 | 28-30 | 1500-1600 |
| Panna | PN | 24.14 | 80.59 | 585.65 | 36-38 | 1200-1300 |
| Rewa | RW | 24.35 | 81.36 | 521.50 | 36-38 | 1200-1300 |
| Sheopur | SP | 25.57 | 76.92 | 356.10 | 36-38 | 800-900 |
| Shivpuri | SHP | 25.44 | 77.72 | 473.00 | 36-38 | 800-900 |

**Fig. 1.**
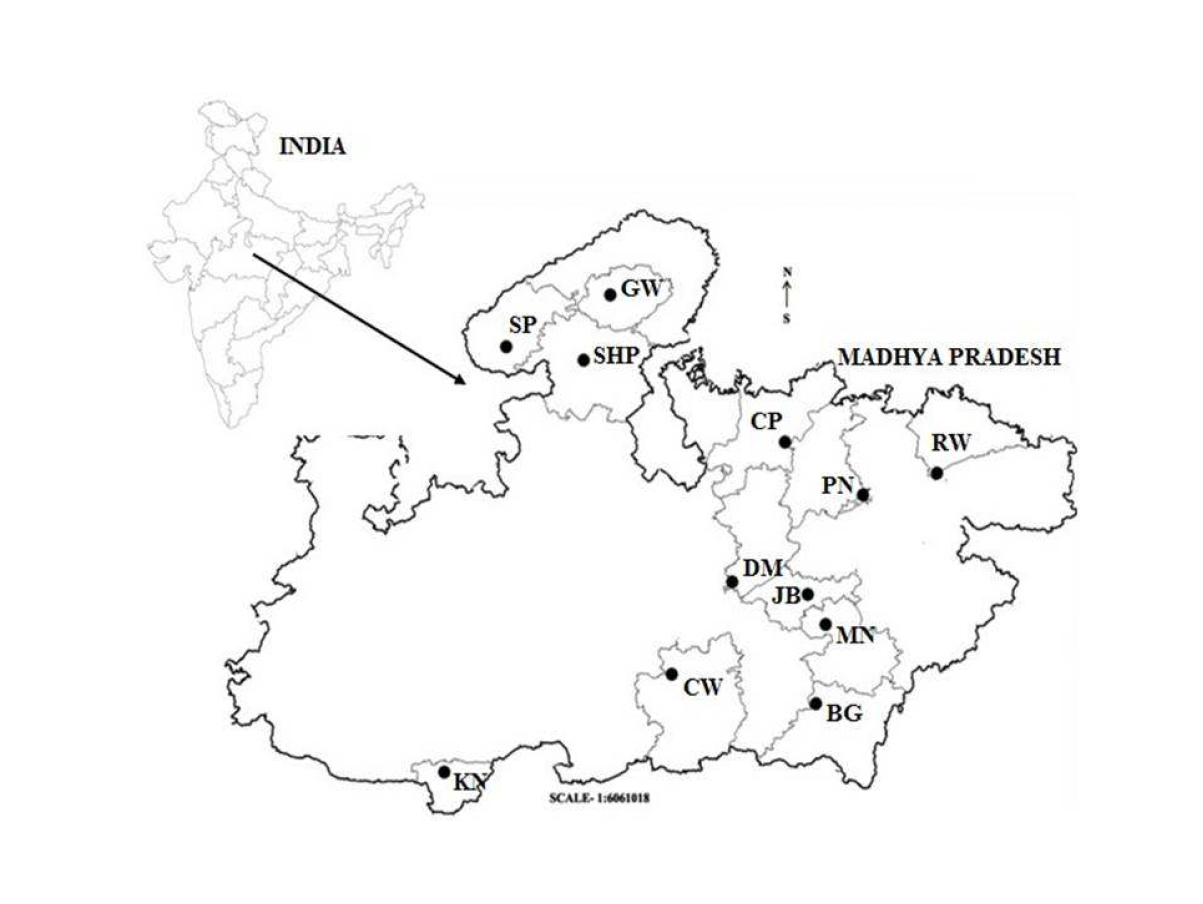
The sampled locations of *B. serrata* metapopulation on the map of Central India (Madhya Pradesh). Details of location codes are given in Table 1

The collected wood core samples were preserved in 40% formaldehyde in plastic tubes. The leaf samples were preserved in cryo-freezer and brought to the laboratory for further assay.

### Measurement of wood fiber length

The wood radial core samples were macerated for separation of wood fibers and to prepare glass slide after staining through 20% safranin following the standardized protocol (Mahesh et al. 2015). Five slides from each of wood core samples were prepared and the length of five fibers per slide were measured using the compound microscope (Leica, Switzerland, Model: EC3) attached with a computer loaded with compatible software LAS version 4.5.0 (Build: 600). Thus, 25 fibers per tree were measured.

### DNA isolation and PCR amplification

DNA from the leaf samples was isolated following modified CTAB method (Deshmukh et al. 2007). 20 dominant primers (10 primers each of RAPD and ISSR) were selected for PCR amplification of all 240 genotypes across all sampled locations (Table 2).

**Table 2.**
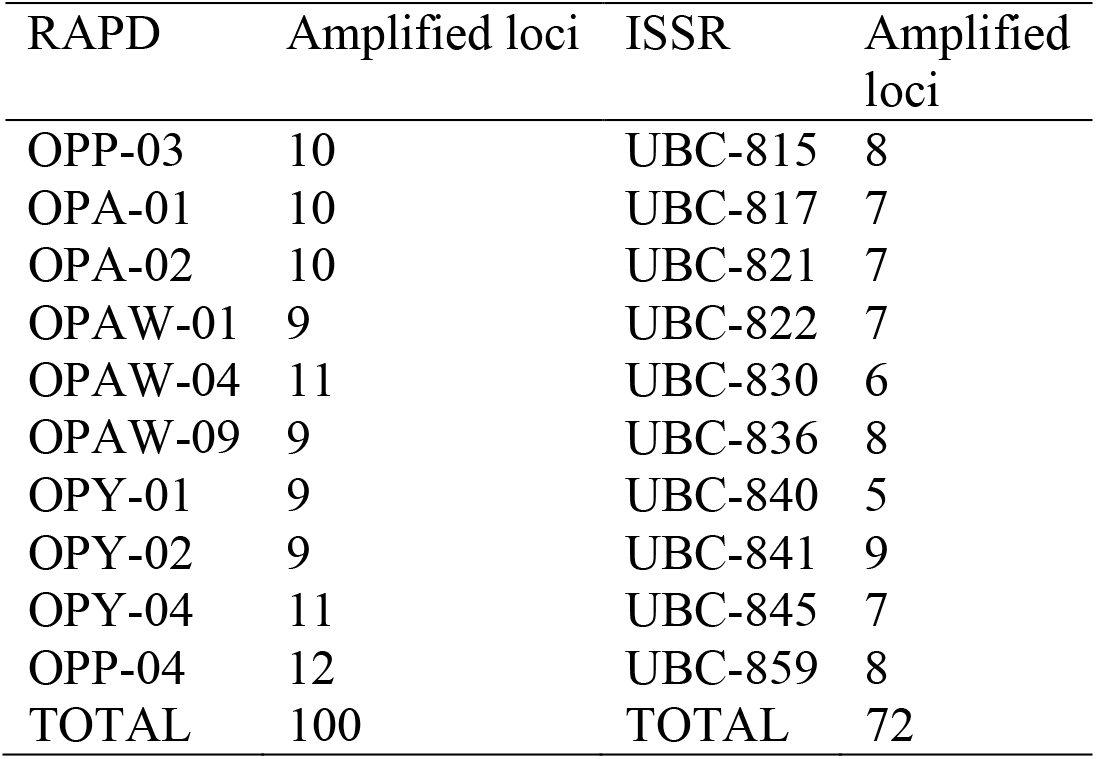
The RAPD and ISSR primers applied for the investigation and their amplification

The reaction mixture for RAPD/ ISSR assay comprised 15/ 10 μl 1x buffer with KCl containing 50 / 40 ng genomic DNA, 0.66/ 0.80 μM primers, 0.2/ 0.4 mM dNTPs and 1.5/ 2.5 μM MgCl_2_ with an uniform 1U Taq polymerase. PCR parameters for RAPD/ ISSR included an initial denaturation steps at 94°C 4min/ 3min followed by 35 cycles for the annealing temperature each at 35 °C/ 50 °C for 30sec, extension each 72°C for 45 sec/ 60 sec followed by final extension at 72 °C for 5 min/ 10 min. The amplified products were electrophoresed on 1.5% agarose gel containing 0.5μgml^-1^ ethidium bromide in 0.5 X TBE (pH 8.0). Separation was carried out by applying a constant voltage of 100V for 3h. The fractionated amplified products on agarose gel were visualized on gel documentation system under UV light. The visualized bands were scored in a Microsoft excel sheet to form a binary data genetic profile.

### Data analysis

Data obtained for fiber length and tree girth were subjected to descriptive statistical analysis including Q-Q and P-P plots for verification of normal distribution in data sets, and analysis of variance, i.e. ANOVA; and if F values found significant at *p* <0.05, LSD at p=0.05, i.e. LSD_0.05_ was computed to compare mean values of both parameters across 12 locations. The statistical analysis was performed using SPSS software program. ANOVA (Table 3) was also used for computation of Q_ST_ (Lande, 1992) values of 66 between paired locations, i.e. Q_ST_ (L) and between populations of the species, i.e. Q_ST_ (P) for fiber length and GBH using the following expressions:

QST =σ^2^GB /(σ^2^GB+2 σ^2^GW)

Q_ST_ (L) = (S_L_-W_L_)/ (S_L_+49W_L_)

Q_ST_ (P) = (S_P_-W_P_)/ (S_P_+39W_P_)

**Table 3.**
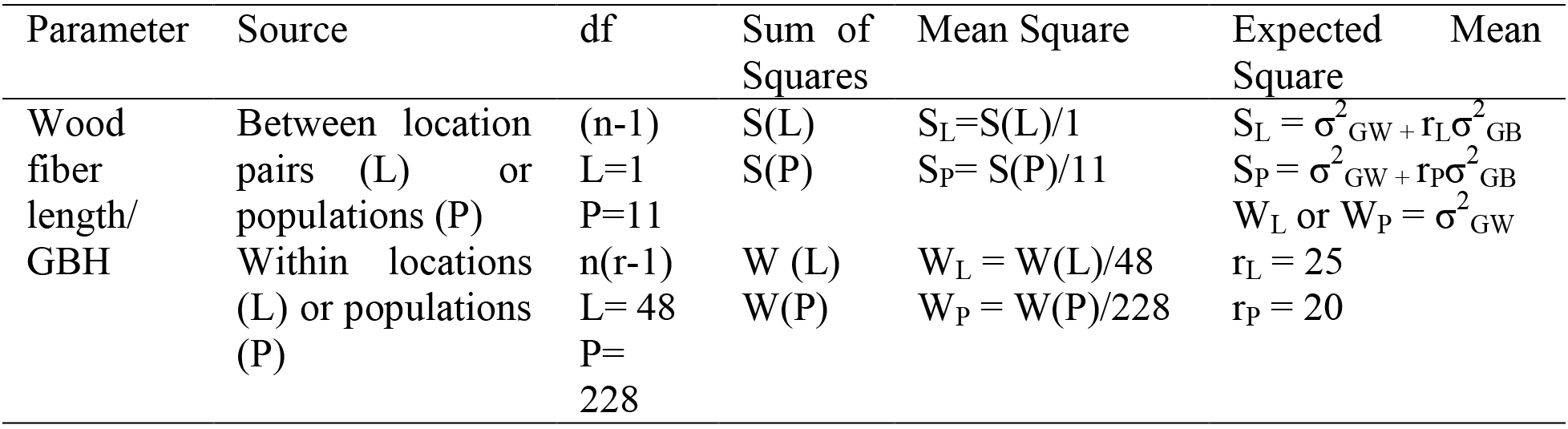
Analysis of variance (ANOVA) model for computation of quantitative trait index (*Q_ST_*) for wood fiber length and GBH of *Boswellia serrata* Roxb.

Φ_ST_ values for 66 paired locations, Φ_ST_ (L) and species, Φ_ST_ (P) were computed from analysis of molecular variance (AMOVA) obtained for neutral molecular marker (RAPD+ISSR) data (Ansari, unpublished), using ARLEQUIN version 3.5 (Excoffier and Lischer, 2010). A comparison between Q_ST_ and Φ_ST_ values was made for ascertaining deviation from null (Q_ST_ = Φ_ST_) hypothesis.

## Result and Discussion

*Boswellia serrata* is an economically important multipurpose tree species and has become endangered in its natural habitat. The genetic worth of a species is assessed in terms of extent of variability existing in morpho-physiological traits of economic importance; the lower the variation range in the traits, the less genetic worth of the species from viewpoint of economics as well as adaptability in the natural habitat. Consequently, the present investigation was undertaken to assess types of selection pressure and/or genetic drift acting on endangered *B. serrata* metapopulation on the basis of quantitative genetic divergence (Q_ST_) with neutral genetic divergence (Φ_ST_).

The data obtained for wood fiber length and GBH was subjected to descriptive statistics in order to attest normal distribution, robustness and outliers. The departure from normal distribution affects tests and confidence intervals based on homogeneity of the data, especially in case of discontinuous populations of endangered species. Table 4 depicts descriptive statistics, which exhibited variations of 59.7% for wood fiber length and 505.2% for tree girth differences between minimum and maximum values across all 12 locations.

**Table 4.** Descriptive statistics for wood fiber length and girth across twelve collected locations of *Boswellia serrata* Roxb.

| Parameter | N | Range | Min | Max | Mean ±SD | Variance | Kurtosis |
| --- | --- | --- | --- | --- | --- | --- | --- |
| WFL (μm) | 240 | 491 | 822 | 1313 | 1042 ±96 | 9175 | -0.344 |
| Girth (cm) | 240 | 192 | 38 | 230 | 105 ±33 | 1125 | 0.697 |

WFL= wood fiber length, N= number of trees, Min= minimum, Max= maximum, SD= standard deviation,

It may be emphasized that the natural metapopulation of *B. serrata* represents large pools of mature trees with variable age groups vis-à-vis wide range of tree girths. However, wood fiber length increases during juvenile phase (young age) and levels off during mature phase (reproductive age) in most tree species (Dinwoodie 1961; Bendtsen, 1978; Bendtsen and Sneft 1986; Zobel and van Buijtenen 1989; Peszlen 1994; Gartner et al. 1997; Adamopoulos et al. 2010). Further, wood fiber length is the least influenced morpho-metric parameter by climatic conditions in comparison to tree growth (Watt et al. 2008). This is the reason that the observed variation in wood fiber length is far less than that in GBH across all sampled trees of twelve natural locations. Hence, wood fiber length appears to be more reliable than GBH for population genetics related investigations because the variation in tree girth confounds the effects of age and environment.

The wood fiber length and tree girth had slight positive and negative kurtosis, respectively (Table 4), which is an estimate of departure from normal distribution and may influence tests of variance and covariance (Jobson 1991, p.55; Mardia et al. 1979, p. 149). Browne (1982, 1984) has also demonstrated that kurtosis largely influences significance tests and standard errors of parameter estimates. The existence of slight kurtosis in the present investigation perhaps demonstrates uneven loss (removal) of genotypes or setting up of genetic drift due to discontinuous and small size of metapopulation of *B. serrata*. To ascertain the normal distribution, the data were subjected to Q-Q plot (non-cumulative probability distribution) and P-P plot (cumulative probability distribution), which magnify deviation from the proposed normal distribution on tails and middle, respectively.

In the present investigation, the data accumulation around straight lines in both Q-Q and P-P plots (Fig. 2) demonstrates a normal distribution trend with insignificant kurtosis, which is unlikely to influence analysis of variance. Thus, the collected data is adequately robust to withstand moderate departures from normal distribution (Harwell et al. 1992; Lindman 1992; Stevens 1996). Further, the large size of samples as in the present case nullifies the impact of kurtosis, if any, on various statistical analysis and test of significance (Tiku et al. 1986). The collected locations exhibited significant variations in wood fiber length and GBH (Fig. 3). Wood fiber length was maximum in Khandwa (KN) location and minimum in Jabalpur (JB) location. However, the wood fiber length obtained in Mandla (MN) and Shivpuri (SHP) locations was on par with that in KN location. The wood fiber length of KN location was 18.2% more than that in JB location. As for GBH, Chhindwara (CW) and Damoh (DM) locations had significantly maximum and minimum values, respectively. However, the GBH of Chhatarpur (CP), JB and Sheopur (SP) locations was on par with that of CW location. GBH of CW location was 109.2% more than that of DM location. Variation in wood fiber length with respect to different sites has been reported in case of *Gmelina arborea* (Moya and Fo 2008) and *Pinus radiata* (Watt et al. 2008). However, there is less information available on variation in wood fiber length across natural populations of a species, except *Dipterocarpus indicus,* wherein the eight natural populations in the Western Ghat have shown significant differences in wood fiber traits (Devi Prasad et al. 2012). The tree/clonal differences for wood fiber length have also been documented in cultivated *Eucalyptus globulus* (Jorge et al. 2000; Wimmer et al. 2008), *Populus* sp. (DeBell et al. 2002), *Populus deltoides* (Pande et al. 2012).

**Fig. 2.**
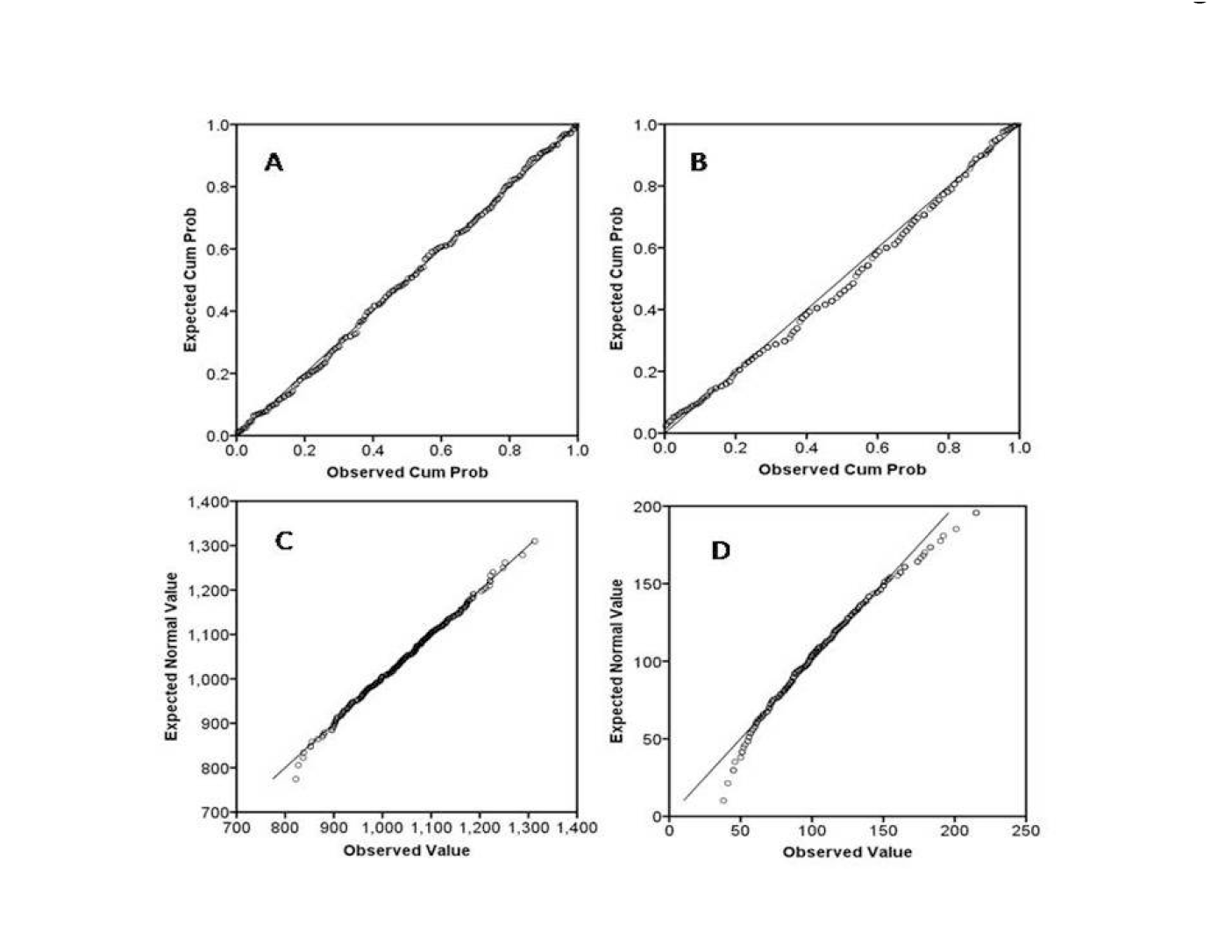
Q-Q Plot (A-B) and P-P plot (C-D) based on non-cumulative probability distribution and cumulative probability distribution, respectively for verification of normal distribution pattern in data for wood fiber length (A,C) and tree girth (B,D)

**Fig. 3.**
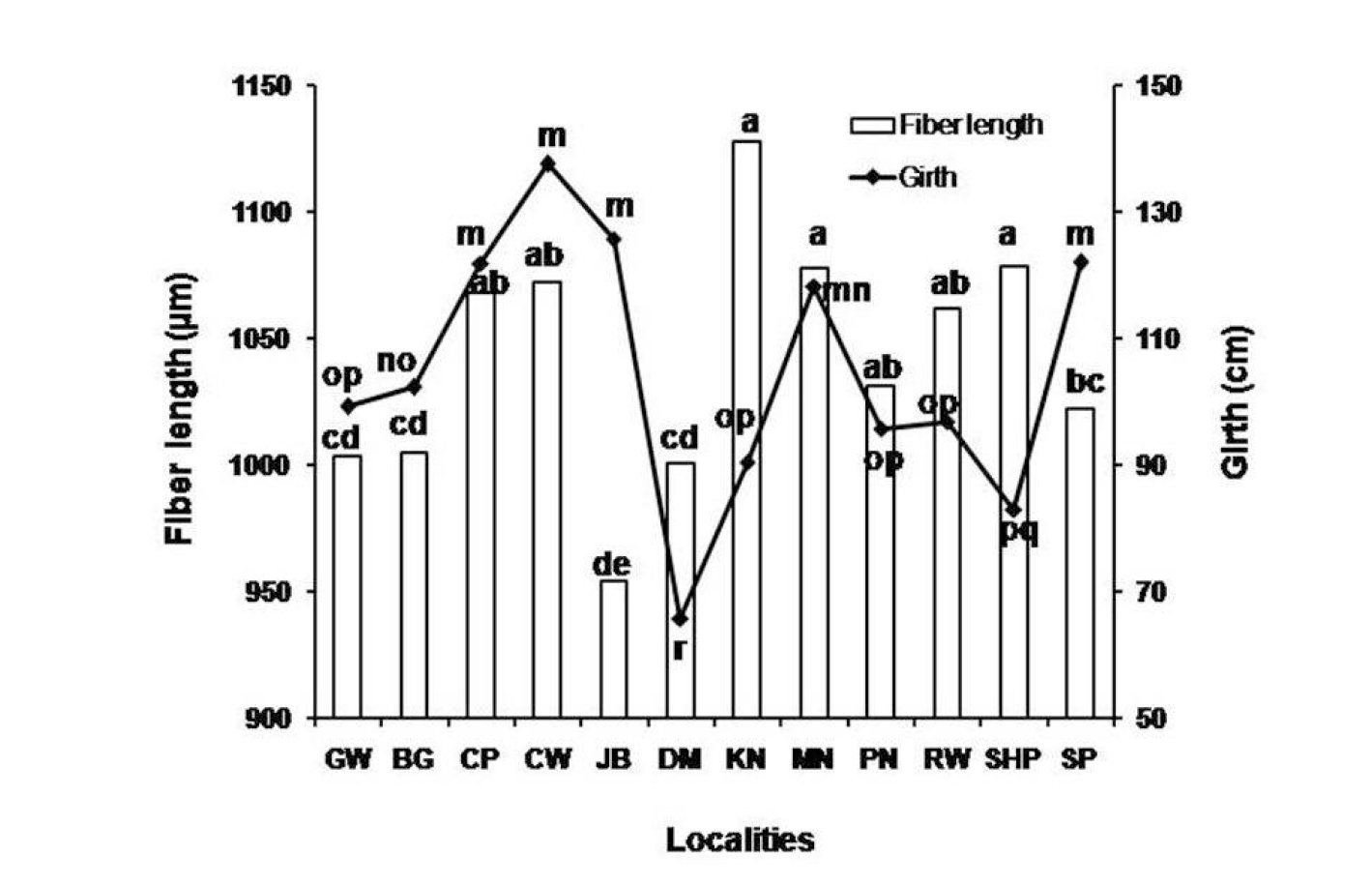
Variation in wood fiber length and tree girth across twelve locations of *Boswellia serrata* Roxb. The points for both parameters exhibiting different letters are significantly different from each other at p < 0.001. Detail of population codes is same as given in Table 1

Thus, *B. serrata* metapopulation is highly variable with respect to GBH compared to wood fiber length. In addition to variable age of trees as mentioned above, the GBH is a manifestation of radial growth, which is influenced by external factors, e.g. temperature, rainfall, soil fertility, inter-/intra-specific competition, pest and diseases, etc. (Watt et al. 2008). The locations with a large GBH in the present investigation mostly belong to relatively moist and moderately warm tropical conditions. In contrast, the locations with better wood fiber length mostly belong to dry and warm conditions. There is no information about climatic conditions responsible for wood fiber formation in tropical trees. However, Rossi et al. (2008)have elaborately demonstrated that the rise in temperature accelerates the process of xylogenesis in temperate conifers. Similarly, Watt et al. (2008)have also inferred that the average air temperature is a major determinant for wood fiber length in *Pinus radiata*. The interpretation also implies independence of the process of wood fiber formation from that of radial (girth) growth; and gains support from the work of Bendtsen and Senft (1986)and DeBell et al. (1998), who have observed no relationship between wood fiber length and growth rate within annual ring in cottonwood, loblolly pine and *Populus* hybrid. However, Koubaa et al. (1998)have reported a slight negative correlation between wood fiber length and growth rate in trees of *Populus* hybrid clones.

We assume that the sampled locations of *B. serrata* in the present investigation are parts of a continuous large metapopulation, which became fragmented due to selective disproportionate removal of superior genotypes, i.e. heterozygotes for commercial exploitation either for manufacturing of paper and pulp or tapping of salai oleoresin gum. The selective removal would have resulted in fragmentation of the metapopulation or creation of population islands. A comparison between Q_ST_ and Φ_ST_ is expected to discern both genetic drift and natural selection process in fragmented populations of *Boswellia serrata*. However, a precise estimation of Q_ST_ needs the data exhibiting normal distribution derived from large number of populations in a balanced design (O’hara and Merila 2005). The data for Q_ST_ estimation in the present investigation conform the criteria of normal distribution as verified by Q-Q and P-P plots (Fig. 2) and a balanced design of 20 trees each of 12 locations. In addition, Q_ST_ estimates are usually made for life-history traits (Morgan et al. 2001, McKay and Latta 2002) from well designed common garden experiments (see review by Villemereuil et al. 2016). Laying out garden experiments is challenging in case of endangered wild species with abortiveness and poor germination of seeds. In a review, Leinonen et al. (2008)have, however, demonstrated that morphological and life-history traits exhibit similar average difference between Q_ST_ and F_ST_; and Q_ST_ estimates from the wild phenotypes to those from the common garden experiment.

The stabilizing selection ascertained by Q_ST_ (L) < Φ_ST_ (L) was observed in 64% paired locations for fiber length and 44% paired locations for GBH. The divergence/ selection trend ascertained by Q_ST_ (L) > Φ_ST_ (L) was observed in 23% paired locations for fiber length and 47% paired locations for DBH. The genetic drift trend ascertained by Q_ST_ (L) = Φ_ST_ (L) was observed in 14 % paired locations for fiber length and 9% paired locations for GBH (Fig. 4). The situation indicates fragmentation of *B. serrata* metapopulation at most locations due to deficit in Q_ST_ (L) values, especially for fiber length, compared to the reference neutral Φ_ST_ (L). The above trend highlights that the sizable paired locations with Q_ST_ (L) < Φ_ST_ (L) exhibit predominance of resilient phenotypes and are in stabilizing act between genetic drift and migration.

**Fig. 4.**
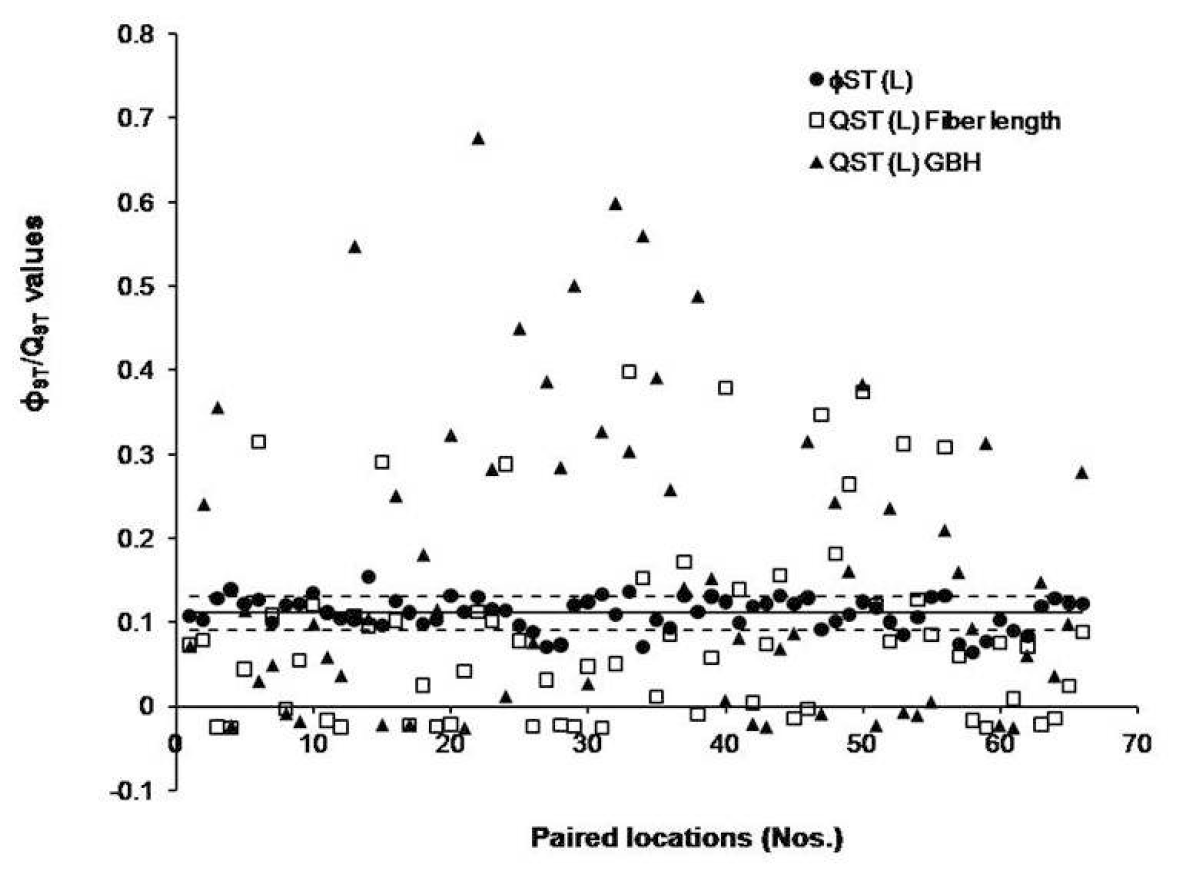
66 paired location wise Q_ST_ (L) estimates for fiber length and DBH. Solid line depicts average Φ_ST_ (L) estimate obtained from RAPD+ISSR molecular marker data and dotted lines for standard deviations for solid line. Q_ST_ (L) > Φ_ST_ (L) is above to dotted line, Q_ST_ (L) = Φ_ST_ (L) between both dotted lines and Q_ST_ (L) < Φ_ST_ (L) below to dotted line of standard deviation

A very few paired locations are also suffering from fragmentation and genetic drift. Both groups comprising bulk of paired locations do not qualify for adaptability and differentiation due to selection and are, hence, dwindling in their habitat. They need to be enriched with divergent genotypes from locations exhibiting Q_ST_ (L) > Φ_ST_ (L). These trends are not discernible from average differentiation analysis of *B. serrata* metapopulation where a singular trend of Q_ST_ (P) > Φ_ST_ (P) has emerged (Table 5). Therefore, the overall comparison between both indices is not much informative and may be misleading as in the present investigation. Clearly, *B. serrata* metapopulation is deficit in requisite genetic diversity at most locations.

**Table 5.**
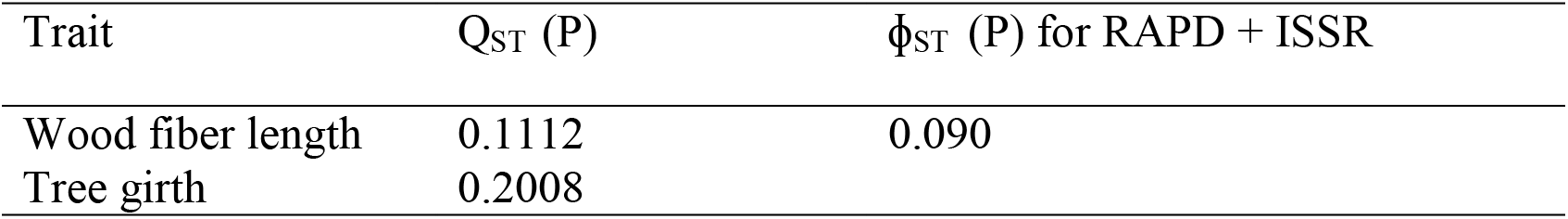
Between population values of quantitative trait index, Q_ST_ (P) and fixation index, F_ST_ (P) for wood fiber length and tree girth in *B. serrata.*

| Trait | $Q_{ST}$ (P) | $\phi_{ST}$ (P) for RAPD + ISSR |
| --- | --- | --- |
| Wood fiber length | 0.1112 | 0.090 |
| Tree girth | 0.2008 |  |

By and large, the location-wise comparison of both indices for metapopulation of a species in the literature is missing. On the other hand, there exists a rich body of literature that has shown comparison of both indices at species level; and in most cases, the Q_ST_ > F_ST_ situation has only been observed (e.g. Merila and Crnokrak 2001; Petit et al. 2001; Edelaar et al. 2008; Ye et al. 2013; Dubois and Cheptou 2016). Further, there is no correlation between pair wise Q_ST_ (L) and pair wise Φ_ST_ (L) for fiber length and GBH. Castillo et al. (2015) have also not found a significant correlation between pair wise P_ST_ (Q_ST_ analogue) and pair wise F_ST_ for three resistance traits under varying scenarios of heritability in *D. stromonium*. It may be reiterated that unlike neutral molecular marker traits, the quantitative traits experience a very strong selection pressure under different geo-climatic conditions and become much more variable and differentiated than the former. As a result, Q_ST_ estimates widely fluctuate in comparison to F_ST_ estimates for the same pair of populations; and thereby no significant correlation between both indices.

## Conclusion

To sum up, *B. serrata* metapopulation exhibits significant variations with respect to wood fiber length and tree girth. Of both parameters, the wood fiber length is less variable across locations. Our investigation reveals that comparison of Q_ST_ (L) and Φ_ST_ (L) across 66 paired locations resolves the status of *B. serrata* metapopulation better than the average differentiation at sub-population level. The analysis brings out three groups, i.e. Q_ST_ (L) = Φ_ST_ (L), Q_ST_ (L) < Φ_ST_ (L) and Q_ST_ (L) > Φ_ST_ (L). The former two groups are in preponderance and exhibit stabilizing selection or genetic drift, pulling the metapopulation towards instability and genetic erosion. Thus, there is a need to enrich *B. serrata* metapopulation with genetic variability, which would facilitate out crossing, seed vigor and viability, natural regeneration and population divergence. The task may be achieved by introduction of genotypes from the genetically well differentiated locations, i.e. Q_ST_ (L) > Φ_ST_ (L). This would create genetic variability, check population fragmentation and arrest genetic drift in order to rescue the species from endangered status.

## Acknowledgements

Financial support received from the Indian Council of Forestry Research and Education, Dehradun under the project ID-175/TFRI/2011/Gen-4 (23) is gratefully acknowledged.

## Compliance with Ethical Standards

The authors declare that they have no conflict of interest.

